# Cerebellar control of targeted tongue movements

**DOI:** 10.1101/2024.09.26.615128

**Authors:** Lorenzo Bina, Camilla Ciapponi, Si-yang Yu, Xiang Wang, Laurens W.J. Bosman, Chris I. De Zeeuw

## Abstract

The cerebellum is critical for coordinating movements related to eating, drinking and swallowing. Cerebellar Purkinje cell activity has been shown to encode ongoing tongue movements, but it is unclear how this activity can alter the trajectory of the tongue. To elucidate the impact of Purkinje cells on goal-directed tongue movements, we recorded their activity in the vermis and hemispheres during spontaneous licking from a stationary or moving water spout. Some Purkinje cells encode rhythmic tongue movements with their complex spikes, others with their simple spikes or a combination of both. Complex spikes predominantly marked the start and end of a licking bout, and thus encoded behavioural state changes, while simple spike firing was more related to individual licks. In addition, complex spikes reported unexpected changes in the position of the water spout and subsequent modulation of simple spike firing caused bending of the tongue, reaching out for the new target position. Using machine learning, we demonstrated that it is possible to predict licking activity based on the spiking patterns of individual Purkinje cells. Using optogenetic stimulation of Purkinje cells, we could experimentally replicate the impact of modulated simple spike firing, suggesting that increased simple spike activity indeed causes ipsilateral bending of the tongue during goal-directed movements. Our data highlight that directional control of movements is paramount in cerebellar function and that complex spike and simple spike modulation complement each other during sensorimotor coordination. These results bring us closer to understanding clinical implications of cerebellar disorders during eating, drinking and swallowing.

**Key points:**

- When drinking, mice make rhythmic tongue movements directed towards the water source.
- Cerebellar Purkinje cells can fire rhythmically in tune with the tongue movements.
- Purkinje cells encode changes in the position of the water source with complex spikes.
- Purkinje cell simple spike firing affects the direction of tongue movements.
- Purkinje cells that report changes in the position of the target can also adjust movements in the right direction.

## Introduction

Cortical and subcortical structures, like the cerebellum and basal ganglia, are essential for controlling complex motor behaviours that necessitate the coordinated action and precise sequencing of multiple muscle groups. This does not only hold for skilled movements of the whole body (Sauerbrei *et al*., 2015; Ting *et al*., 2015; Vinueza Veloz *et al*., 2015; Kim *et al*., 2017), but also for those of individual limbs (Diedrichsen *et al*., 2005; Park *et al*., 2022) or the tongue (Leopold & Kagel, 1996; Travers *et al*., 1997; Cullins *et al*., 2019). Indeed, various neurological conditions can compromise complex motor behaviours, and tongue movements are no exception: dystonia, dyskinesia and/or tremor of the tongue, as well as impaired oral sensation can impair such crucial actions like mastication, drinking, swallowing, breathing or speaking (Leopold & Kagel, 1996; Ghadery *et al*., 2022; Laurence-Chasen *et al*., 2022). Eventually, improper control of the tongue can lead to the life-threatening condition of aspiration pneumonia (Takizawa *et al*., 2016; Krohn *et al*., 2023).

Accordingly, patients suffering from diseases of the cerebellum, such as hereditary cerebellar ataxia caused by malfunction of cerebellar Purkinje cells, can show symptoms like dysarthria, dysphagia or choking due to reduced motor control of the tongue (Ikeda *et al*., 2012; Ushe & Perlmutter, 2012; Markovic *et al*., 2016; Keage *et al*., 2017; Rezende Filho *et al*., 2019; Woo *et al*., 2019; Giardina *et al*., 2020). In line with the latter, functional imaging in human subjects has revealed specific activation patterns in the cerebellar cortex and nuclei during both voluntary and subconscious tongue movements (Corfield *et al*., 1999; Dimitrova *et al*., 2006; Ogura *et al*., 2012; Groenendijk *et al*., 2020; Sörös *et al*., 2020), while electrophysiological recordings in Purkinje cells and other cerebellar neurons during licking have shown close correlations between cerebellar activity and tongue movements (Welsh *et al*., 1995; Bryant *et al*., 2010; Lu *et al*., 2013; Gao *et al*., 2018; Bina *et al*., 2021; Gaffield *et al*., 2022; Lackey *et al*., 2024). However, despite the experimental and clinical evidence for a role of the cerebellum in controlling tongue movements, our understanding of the mechanisms by which neural activity of cerebellar neurons can affect tongue movements remains limited.

The primary driver of tongue movements is a central pattern generator consisting of premotor neurons in the intermediate reticular formation of the brainstem (Brozek *et al*., 1996; Travers *et al*., 1997; Dempsey *et al*., 2021; Kleinfeld *et al*., 2023). These reticular neurons directly activate motor neurons in the hypoglossal nucleus, which control all four intrinsic and three out of the four extrinsic tongue muscles, while they may indirectly also affect the vagal nerve that drives the fourth extrinsic muscle (Wiesenfeld *et al*., 1977; Altschuler *et al*., 1994; Guo *et al*., 2020). As a consequence, a lack of the cerebellar output that influences the activity of the reticular neurons reduces the efficiency of tongue movements (Bryant *et al*., 2010), while it does not preclude making tongue movements per se (Timmann *et al*., 2003). Given the multitude of muscles required for precise tongue movements, we hypothesize that the cerebellum is required for the coordination of these muscles, similar to its function of inter-limb coordination to allow walking over a horizontal ladder (Vinueza Veloz *et al*., 2015; Jaarsma *et al*., 2024).

If the cerebellum is indeed able to coordinate the activation of the tongue muscles so that tongue movements can be adapted to changes in the behavioural needs, then Purkinje cells should encode unperturbed tongue movements, react to environmental changes and translate these into altered movements. Using statistical analysis and a machine learning model, we demonstrate that Purkinje cells indeed encode spontaneous licking. In particular, complex spikes occur during behavioural state changes, thus the start and end of licking bouts, while simple spikes are associated with individual licks. In addition, complex spikes report fast changes in the position of the water spout, and this is correlated to alterations in the simple spike firing, causing a rapid adaptation of the tongue trajectory. Finally, using optogenetic stimulation of Purkinje cells, we could demonstrate the causal impact of simple spike firing on the tongue trajectory. Thus, cerebellar Purkinje cells meet the criteria for mediating adaptation of tongue movements to changes in environmental demands.

## Methods

### Ethical approval

All experimental procedures involving animals were evaluated before the start of this study by an independent animal ethical committee (DEC-Consult, Soest, the Netherlands) and subsequently approved by the national authority (Centrale Commissie Dierproeven, The Hague, The Netherlands; project license AVD1010020197846). Compliance with the project license of each experiment was confirmed by the animal welfare body of the Erasmus MC, and all experiments were performed conform the relevant institutional regulations of the Erasmus MC, Dutch legislation on animal experimentation, and EU directive 2010/63/EU. Throughout the study, care was taken to minimise pain and suffering.

### Mice

All experiments that did not involve optogenetic stimulation were performed with C57BL/6J mice (Charles River, ‘s Hertogenbosch, The Netherlands). For experiments with optogenetic stimulation, Tg(Pcp2-cre)2Mpin; Gt(ROSA)26Sor^tm27.1(CAG-OP4*H134R/tdTomato)Hze^ mice on a C57BL/6J background were used (Witter *et al*., 2013), and these mice were bred in the animal facility of the Erasmus MC with regular back crossings with wild-type C57BL/6J mice. All mice were adults between 12 and 35 weeks of age and specific pathogen free. The animals were group housed until surgery; after which they were single housed to avoid inflicting wounds by cage mates. Mice were kept in a vivarium with controlled temperature and humidity and a 12/12h light/dark cycle.

The mice had ad libitum access to standard food and water at all times, except when they were trained to lick from the water spout in the recording setup. During training, mice had access to water only during the daily sessions, while food remained available ad libitum. If a mouse did not drink sufficiently in the setup, it received the daily dosage of water (1 ml / 20 g body weight) about half an hour after the training session. Daily weighing was used to monitor putative weight loss, which was limited at 20% of the original body weight according to the restrictions in our project license. Typically, however, mice did not lose that much weight: after a first drop of about 10%-15% in the first five days, the body weight typically became stable or even increased once mice got used to drink ad libitum from the setup and their metabolism adapted.

A total of 39 mice were used for electrophysiological recordings, nine of which were trained in the motor adaptation task, and eight mice were used for optogenetic stimulation. We used approximately equal numbers of male and female mice. Given the simplicity of the behavioural task, all mice reached the final stage of the experiments. Only in rare cases when the electrophysiological recordings were not successful, mice were excluded from the animal count. At the end of the final experimental session, mice were euthanized by cervical dislocation under isoflurane anaesthesia.

### Surgical procedures

Mice received a magnetic pedestal for head fixation, attached to the skull above bregma using Optibond adhesive (Kerr Corporation, Orange, CA, USA), and a craniotomy was performed to expose the recording area (the lateral part of vermal lobules VI an VII and the adjacent lobules hemispheric lobules crus 1 and crus 2 on the right side). Four mice received a larger and more central craniotomy in order to expose both right and left vermis for optogenetic stimulation. The craniotomy was cleaned and afterwards covered with Kwik-Cast (World Precision Instruments, Sarasota, FL, USA). Surgeries were performed under isoflurane anaesthesia (induction: 4% V/V; maintenance: 2% V/V in O_2_) with lidocaine (4 mg/kg; Braun, Melsungen, Germany) applied at the surgical locations. Postsurgical pain was treated with carprofen (5 mg/kg s.c.; Pfizer, New York, NY, USA), buprenorphine (50 μg/kg s.c.; Indivior, Richmond, VA, USA), and bupivacaine (1 mg/kg s.c.; Actavis, Parsipanny-Troy Hills, NJ, USA). Three days of recovery followed the procedure.

### Habituation

After recovery from surgery, mice were handled daily by the experimenter to reduce stress. When mice were relaxed when handling, they were head-fixed for ∼15 minutes a day during which water was available from the lick-port positioned in front of the mouse.

### Lick detection

During training and experimental sessions, mice had access to a water spout in front of their mouth. At this water spout, water was freely available. Licking was detected using an optical sensor placed 2 mm before the water spout. In a subset of experiments, tongue trajectories were recorded with an over-head video camera (frame rate 100 Hz). The coordinates of the tip of the tongue were extracted using DeepLabCut software for pose estimation (Mathis *et al*., 2018). We used the location of maximal protrusion for further analysis. From the video analysis it became clear that the moment of maximal protrusion was on average achieved 16 (2) ms after detection by the optical sensor. Licking bouts were defined as sequences of licks with intervals <500 ms.

### Moving lick-port

To test the ability of mice to target their tongue movements, we trained 9 mice to lick from the same lick-port as used in the other experiments, but for these mice, we moved the lick-port 3 mm to the right at unpredictable moments. At the start of each session, the lick-port was in front of the mouse. We used a closed-loop control circuit to detect licking. Once a lick was detected, there was a 50% chance to trigger a rightward movement, starting 40 ms after lick detection and with a travel time of 50 ms. For the movement, we used a piezo actuator (Physik Instrumente, Karlsruhe, Germany). In this way, we moved the lick-port after retraction of the tongue retraction. The lick-port was moved back to the central position after 750 ms and stayed there at least 750 ms before it could be moved again. After a week of training, mice were able to adjust the direction of their tongue movements to follow the lick-port.

### Electrophysiology

We recorded Purkinje cell activity extracellularly in awake mice both during spontaneous licking and motor adaptation, using quartz-coated platinum/tungsten electrodes (R = 2-5 MΩ, outer diameter = 80 μm, Thomas Recording, Giessen, Germany). Electrodes were randomly placed in an 8 × 4 matrix (Thomas Recording), with an inter-electrode distance of 305 μm. Prior to the recordings, the mice were lightly anaesthetized with isoflurane to remove the dura mater, bring them in the setup and place the electrodes on the surface of the cerebellum. Recordings started at least 60 min after termination of anaesthesia and were made at a minimal depth of 500 μm. The voltage signal was digitized at 25 kHz, using a 1-6,000 Hz band-pass filter, 22x pre-amplified and stored using a RZ2 multi-channel workstation (Tucker-Davis Technologies, Alachua, FL, USA). Once awake, the attention of the mice was triggered by randomly delivering a few drops of water until they spontaneously started seeking for water.

Spikes were detected off-line using SpikeTrain (Neurasmus, Rotterdam, the Netherlands). A recording was considered to originate from a single Purkinje cell when it contained both complex spikes (identified by a stereotypic waveform, overshooting and the presence of spikelets) and simple spikes, and in which each complex spike was followed by a pause of at least 8 ms before simple spike firing resumed. We included in our analysis all the recorded cells for which the quality remained constant for a at least 20 trials. When looking at complex spike and simple spike modulations, we built peri-stimulus time histograms (PSTHs) using bins of 10 ms. Modulation depth was defined as the difference between maximum and minimum fluctuation in the average spike modulation to each lick of a bout within a window of 200 ms.

### Phase analysis

We used a linear phase transform as described previously (Romano *et al*., 2020). The phase values of 0, π and 2π correspond to the moments of protrusion start, maximal protrusion, and end of the retraction, respectively. We considered the values of magnitude-squared coherence and cross spectrum phase between spike-lick cross correlograms and the lick autocorrelograms.

### Optogenetic stimulation

After learning to lick fluently from the lick-port, one or two optic fibres (diameter 400 μm, Thorlabs, Newton, NJ, USA) were placed on the brain surface. With the optic fibres, we could apply blue light (470 nm, 5 mW) for optogenetic stimulation. In a group of four mice, we stimulated vermal Purkinje cells left and right of the midline. In four other mice, we stimulated the right paravermis and the adjacent and more lateral part of lobule VI. We gave pulses with a duration of 160 ms (corresponding to approximately one inter-lick interval) starting 10 ms after laser detection of a randomly selected lick. Stimulation of either one or both optic fibres was randomly intermingled. In a third set of experiments, we stimulated only right paravermal Purkinje cells, alternating 160 ms pulses with 80 ms pulses starting 10 ms after lick detection or 80 ms pulses starting 90 ms after lick detection. An extra session was performed with the last group of mice, during which we recorded spikes from Purkinje cells during light stimulation to test its efficacy.

### Model

To develop a model that explains the relationship between Purkinje cell activity and licking behaviour, we adapted an XGBoost classifier model (Chen & Guestrin, 2016). Initially, we processed the raw data on spikes and behaviour, cutting it into 200 ms snippets. These snippets included a 10 ms overlap to ensure continuity. Within these snippets, we calculated three key features: simple spike frequency, simple spike CV2, and complex spike frequency. CV2 was calculated as follows: 2 |ISI_n+1_ -ISI_n_|/(ISI_n+1_ + ISI_n_), where ISI = inter-spike interval (Shin *et al*., 2007). The snippets were then grouped based on observed behaviour (presence or absence of licks during a snippet). To create a balanced training set, we randomly selected the same number of snippets from epochs with and without licking. The number was defined by two-thirds of the total amount of the smaller of the two groups. The remainder of the data served as the test set for evaluating our model. For training, we configured the model with 4096 estimators and set the maximum tree depth to 8. We utilized a faster histogram optimized approximate greedy algorithm for enhancing the gradient boost tree. The performance of the trained models was assessed using the test set to determine the correlation between spike activity and licking behaviour. A cell was deemed to exhibit strong licking-correlated spiking if the model’s testing accuracy exceeded 65%. To elucidate the influence of each feature on the model’s output, we employed a game-theoretic method known as SHAP (SHapley Additive exPlanations) (Lundberg *et al*., 2020). By applying SHAP to our validated models, we were able to quantify the contribution of each feature to the model’s predictions, providing insights into the underlying mechanisms linking neural activity to behaviour. For each cell and each parameter, the SHAP value was calculated as the mean of the absolute values of the SHAP values per 200 ms window.

Purkinje cells were considered to be predictive of licking behaviour, if both the accuracy of predicting licking bouts and that of predicting non-licking were >65%. The Purkinje cells performing over 65% correct for licking and non-licking while having a balanced prediction (<15% difference in accuracy for licking vs. non-licking) were considered strongly correlated to licking.

### Statistical analysis

Z-scores [(value – average) / SD] were calculated using the average and SD of baseline activity. Depending on the analysis, we used different baseline intervals. To calculate spiking modulation around the beginning of licking bouts, the baseline was -1000 to -500 ms before bout onset. For the end of licking bouts, we used as baseline the period from 500 to 1000 ms after bout end. To normalize the spike modulation to licks of different coordinates, we used the period from 1000 to 250 ms prior to the licks of interest.

For analysis of the motor adaptation experiments, licks coordinates were normalized to the coordinates of licks prior to the movement of the lick-port; complex spike modulation to target movements was normalized to a similar 500 ms baseline preceding the beginning of licking bouts as for lick timing modulation; simple spike modulation to licks of interest following target movements was normalized to a similar 750 ms baseline preceding the licks of interest as for the analysis of tongue coordinates during spontaneous licking.

Data were tested for normality using Kolmogorov-Smirnov tests, and when normally distributed paired t-tests were used, unless indicated otherwise. Unless stated afterwards, data were summarized as averages (SD).

## Results

Purkinje cells can encode different forms of rhythmic behaviour, like walking, breathing and whisking (Sauerbrei *et al*., 2015; Romano *et al*., 2018; Romano *et al*., 2020). Even though the complex spike activity and the simple spike activity of Purkinje cells have both been shown to correlate with rhythmic licking in separate experiments (Welsh *et al*., 1995; Bryant *et al*., 2010; Cao *et al*., 2012; Gaffield *et al*., 2022), it has remained unclear to what extent the combined activity of complex spikes and simple spikes of individual Purkinje cells can encode licking. Given that the interaction between complex spike and simple spike activity is essential for cerebellar learning, many studies on cerebellar function define Purkinje cells as task-related when they display both complex spike and simple spike modulation during a specific behaviour (Simpson & Alley, 1974; Ito, 2000; Yang & Lisberger, 2014; Ten Brinke *et al*., 2015; Sedaghat-Nejad *et al*., 2022).

### Purkinje cells relate to spontaneous licking

To study putative interactions between complex spike and simple spike firing during rhythmic licking, we accustomed mice to lick from a water spout in front of their mouth, while being head-fixed in the recording setup. Under these conditions, mice licked rhythmically in bouts with lick frequencies generally between 4 and 9 Hz (Fig. 1A), similar to what can be observed in freely moving mice (Boughter *et al*., 2007). During sessions with spontaneous licking, we recorded the activity of Purkinje cells in lobules VI and VII of the vermis and the adjacent hemispheric lobules crus 1 and crus 2. In many, but not all Purkinje cells, we observed rhythmic modulation of complex spike and simple spike activity along with rhythmic licking. More specifically, when we analysed the recordings of the entire periods of licking, in 21 out of 84 (25%) recorded Purkinje cells, complex spike modulation was considered statistically significant (Z > 3), while statistically significant simple spike modulation occurred in 58 out of 84 (69%) Purkinje cells. Notably, only 19 (23%) of the recorded Purkinje cells showed modulation of both their complex spikes and their simple spikes (Fig. 1B-D).

**Fig. 1.**
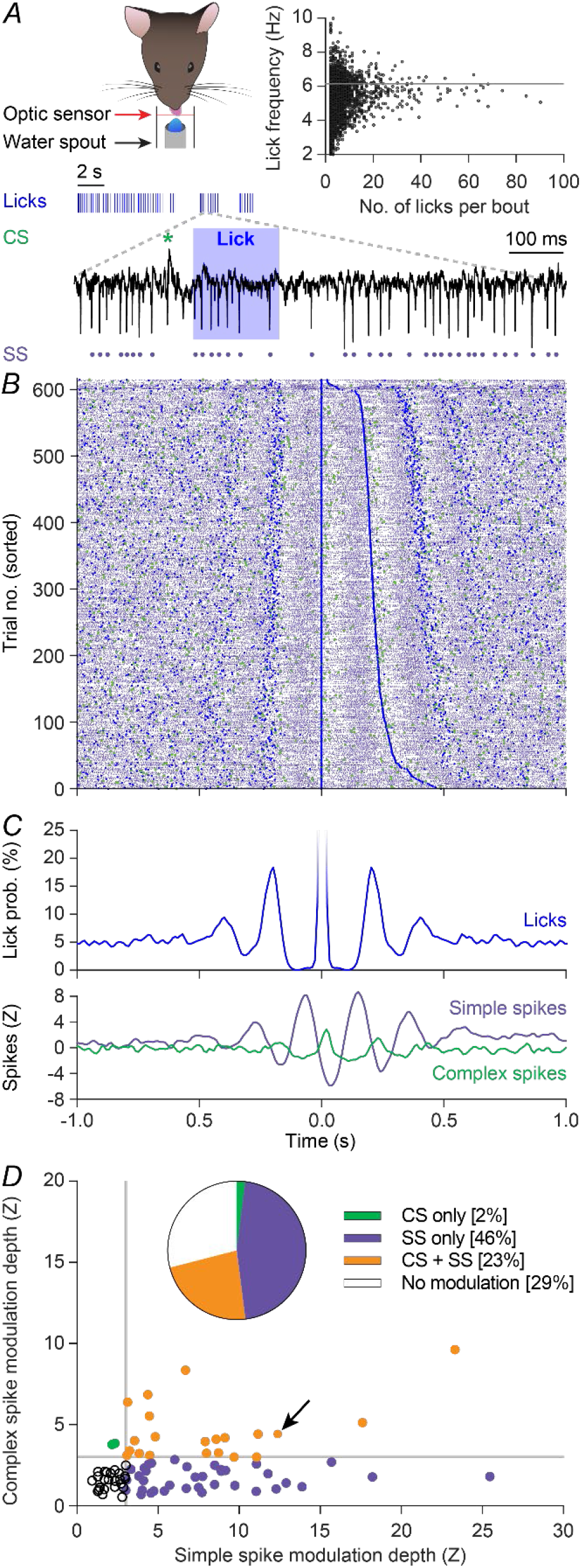
Purkinje cells encode rhythmic licking. **A -** Mice make repetitive tongue movements when licking, and these licks are organized in bouts. In the experimental setup, the median lick frequency of all bouts was just above 6 Hz (horizontal line in inset). To correlate tongue movements to activity of cerebellar Purkinje cells, we made extracellular recordings revealing the timing of complex spikes (green symbol) and simple spikes (purple symbols). **B** - Raster plots showing the distribution of licks (blue), complex spikes (green), and simple spikes (purple) during one session with spontaneous licking. Trials are sorted based on the duration of the inter-lick-interval. **C** - Peri-stimulus-time-histograms (PSTHs) from the licks (top) and spikes (bottom) from the session shown in B. Notice that all three PSTHs display oscillations at similar frequencies. **D** - In individual Purkinje cells, the modulation depths of complex spikes and simple spikes were not necessarily correlated, with most Purkinje cells showing a stronger modulation of their simple spikes than of their complex spikes. Data from 84 Purkinje cells in 32 mice. The arrow indicates the example cell shown in A-C.

*Complex spikes signal behavioural state changes* In view of this surprisingly low number of Purkinje cells displaying both complex spike and simple spike modulation during rhythmic licking, we wondered whether we might have missed specific contributions of complex spikes. As complex spike firing has been associated with changes in behavioural states (Wagner *et al*., 2021), we next focussed on the start and end of licking bouts. In the 300 ms interval centred on the detection of the first lick of a bout, we observed a statistically significant increase (Z > 3) of complex spike firing in 33 out of 84 (39%) recorded Purkinje cells. Similarly, the end of a licking bout corresponded with increased complex spike firing in 43 (51%) Purkinje cells (Fig. 2A-D). Although more Purkinje cells were active around the end than around the start of a licking bout, this difference was not statistically significant (p = 0.163, Fisher’s exact test). A substantial part of the Purkinje cells with significant changes in complex spike firing around state changes signalled both the start and the end of licking bouts (n = 18 [30%]); the others signalled either the start or the end (Fig. 2E, inset).

**Fig. 2.**
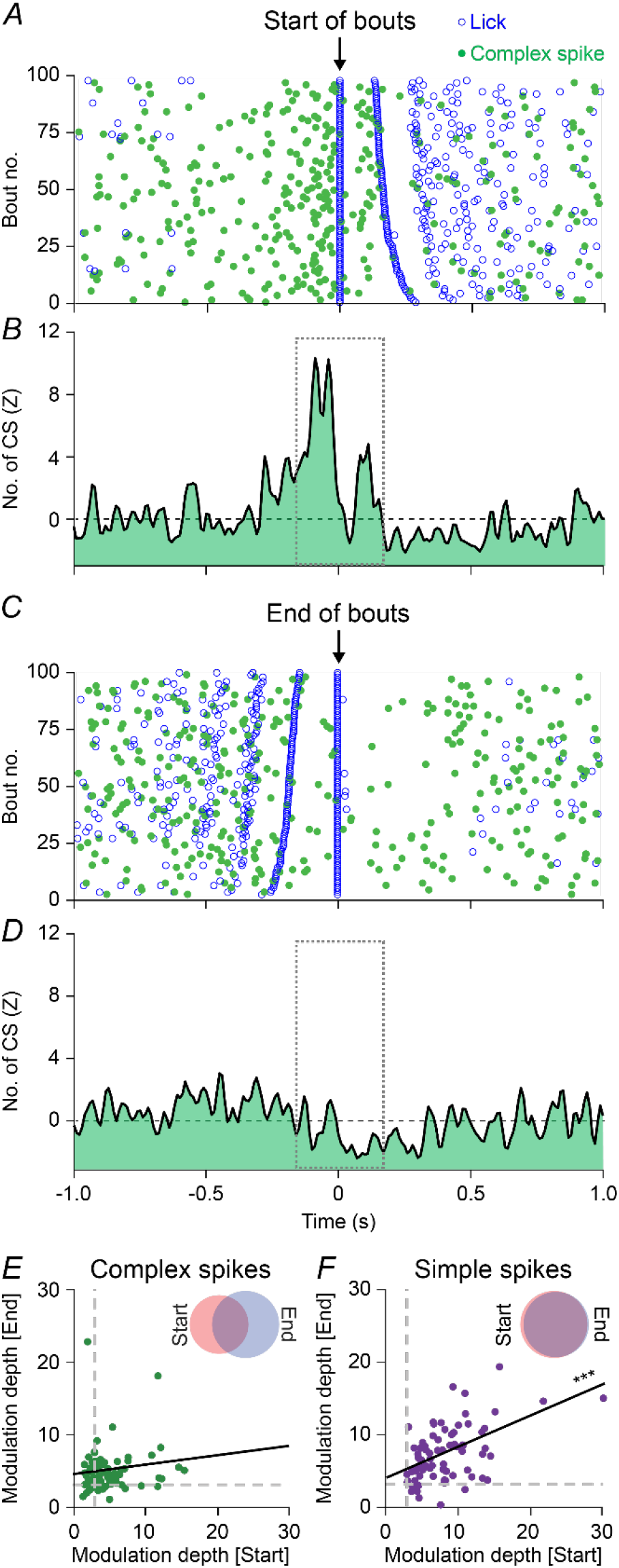
Complex spikes signal behavioural state changes. **A** - The distribution of licks and complex spikes around the start of licking bouts during one session with spontaneous licking. The bouts are sorted based on the duration of the first inter-lick interval. Panels A-D are from the same recording session. **B** - Complex spike PSTH around the start of licking bouts. **C** - The distribution of licks and complex spikes around the end of licking bouts, with the bouts sorted on the duration of the last inter-lick interval. **D** - Complex spike PSTH around the end of licking bouts. **E** - The modulation depths of complex spike firing around the start vs. the end of licking bouts showed no statistically significant correlation (r = 0.015, p = 0.894, n = 84 Purkinje cells, Spearman correlation). Notice that in 69% of the recorded cells the complex spike modulation depth is larger than 3 SD, while only 25% exceeded the threshold for complex spike firing rate modulation around individual licks (Fig. 1D). **F** - Same as E, but for simple spike modulation depth. All recorded cells modulated significantly around either the start or end of licking bouts, and the modulation depths around the start and end of licking bouts were highly correlated per Purkinje cell (r = 0.538, p < 0.001, n = 84 Purkinje cells, Spearman rank correlation).

The notion that the Purkinje cells signalling the start of a licking bout are not necessarily the same as those signalling the end of a licking bout was underscored by the lack of a correlation between the amplitudes of complex spike modulation between both state changes (r = 0.015, p = 0.894, n = 84 Purkinje cells, Spearman correlation, Fig. 2E).

Simple spike modulation, in contrast, was more related to single licks (Fig. 1), and as a consequence simple spike modulation around the first and last lick of a bout were strongly correlated (r = 0.538, p < 0.001, n = 84 Purkinje cells, Spearman correlation, Fig. 2F). Thus, complex spike firing was more related to changes in the behavioural state, i.e. starting or ending a licking bout, while simple spike firing was more related to the execution of individual licks.

### Decoding of Purkinje cell activity patterns with machine learning inference

To investigate whether the observed alterations in Purkinje cell spiking patterns are specific for licking behaviour or whether they also occur during periods without licking, we implemented a machine learning workflow. First, using a 200 ms sliding window, we computed for each recorded Purkinje cell three key features: complex spike frequency, simple spike frequency and simple spike irregularity (CV2). These features were then grouped, based on the presence or absence of licking during each of the 200 ms intervals. An adapted version of the XGBoost classifier model (Chen & Guestrin, 2016) was trained for each cell to detect the relationship between cell activity and licking behaviour. This decoding model identified the activity of 58 out of the 84 (69%) Purkinje cells bearing at least some relation to licking behaviour. Of these Purkinje cells, 28 (i.e., 33%) showed a strong correlation. In general, our model had a similar performance when predicting bout vs. inter-bout intervals, but was slightly better in predicting bout intervals in the cells with the stronger correlations (for all 84 Purkinje cells: 64% for bouts and 64% for inter-bout intervals, p = 0.912, t = 0.112; for the 28 Purkinje cells with the strongest correlations: 76% vs. 73%, p = 0.0471, t = 2.080, paired t tests).

To understand which of the three key features (complex spike frequency, simple spike frequency or simple spike CV2) had the strongest predictive value, we employed the SHAP method on the 58 Purkinje cells that showed a statistically significant simple spike modulation related to licking (Lundberg *et al*., 2020). In line with this selection, it turned out that the absence or presence of complex spike firing of these cells was a poor predictor for being in a licking bout or not. This is visualized for two example Purkinje cells, where the SHAP values dwell around 0, implying weak predictive power (Fig. 3A, B). This does not imply that complex spikes are not related to timing of individual licks, but that they were generally not more present or absent during periods with or without licking in this group of cells. In some Purkinje cells, changes in simple spike firing rate were predictive of licking, whereas in others, changes in simple spike regularity were more predictive, while yet other Purkinje cells displayed changes in both rate and regularity. In the Purkinje cell illustrated in Fig. 3A, the SHAP values of both the simple spike frequency and CV2 were centred around zero, although less so than those of the complex spikes, indicating moderate predictive abilities. However, in the second Purkinje cell (Fig. 3B), the values of the simple spike CV2 were highly asymmetric, with relatively high values (irregular firing) during licking bouts. The absolute SHAP values of simple spike frequency and regularity were unrelated (r = 0.0561, p = 0.676, Spearman correlation, Fig. 3C), indicating that rate and pattern encodings were relatively evenly distributed over Purkinje cells. Overall, however, simple spike firing was a much more accurate predictor of licking behaviour than complex spike firing (p < 0.001, Friedman’s ANOVA, Fig. 3D). Thus, especially alterations in simple spike activity could be good predictors of whether a mouse is licking or not.

**Fig. 3.**
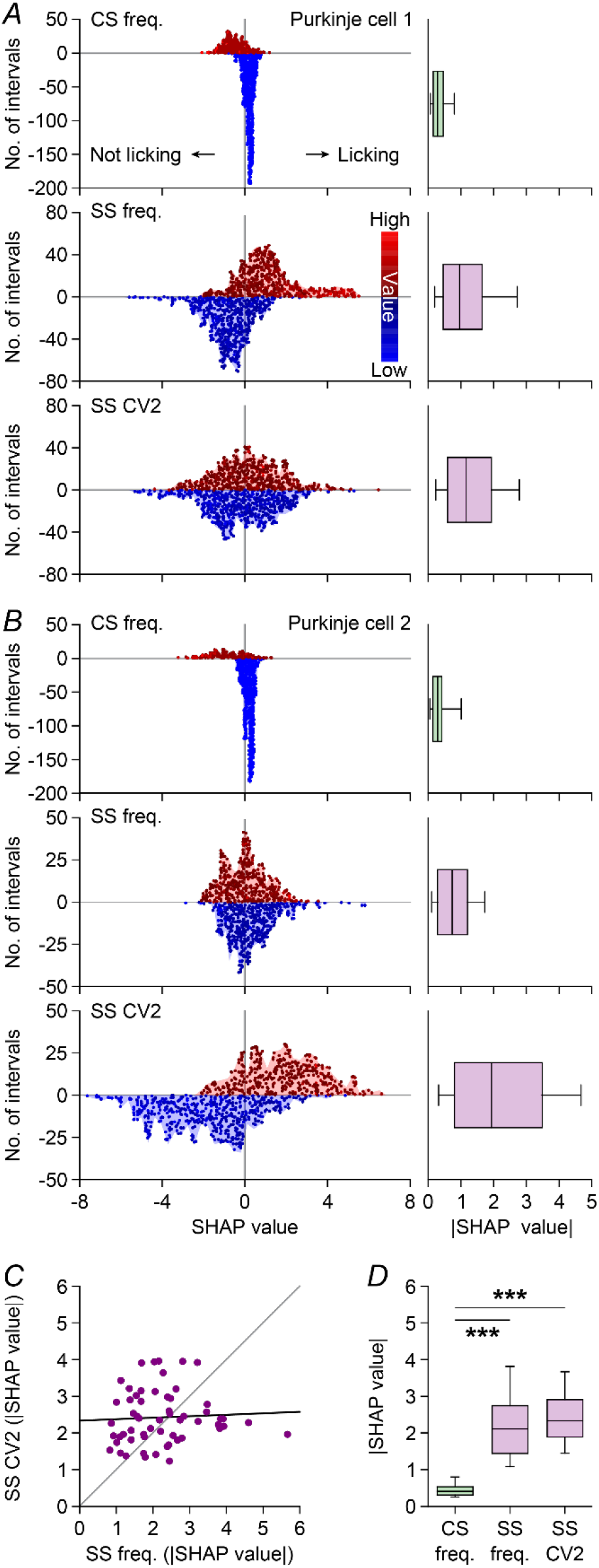
Decoding of Purkinje cell activity patterns with machine learning inference. **A -** We adapted an XGBoost classifier model that could explain the relationship between Purkinje cell activity and licking behaviour. Our model based its prediction on three features extracted from the dataset after dividing it in sliding windows of 200 ms (with 10 ms overlap): complex spike frequency, simple spike frequency, and simple spike CV2. These features are colour-coded based on their values, and for clarity low values (blue shades) are plotted negatively and high values (red shades) positively. Each snippet is plotted individually, and the blue/red contours display a histogram with the distribution of the snippets. For this example Purkinje cell, both simple spike frequency and CV2 are relevant, as illustrated by the bar plots on the right. **B** - Same as in A, but for a different Purkinje cell for which simple spike CV2 has a higher predicting value than simple spike frequency. **C** - The absolute SHAP values for the simple spike related features in the 58 lick-related Purkinje cells; these did not show any correlation (r = 0.056, p = 0.676, Spearman rank correlation). **D** - Absolute SHAP values for the three features reveal much lower impact of the complex spike frequency for the model prediction (p < 0.001, Friedman’s ANOVA, with post-hoc tests vs. complex spikes both p < 0.001).

### Complex spikes and simple spikes cover whole of the lick cycle

Given that our previous analyses confirmed modulation of complex spike as well as simple spikes during licking, we subsequently studied the phase relationships between Purkinje cell activity and licking. To this end, we made a phase transform of all spike times and subsequently performed, for each Purkinje cell, a coherence analysis between licking and spiking. As the phase transform annihilated timing differences between individual licks, this phase-based coherence analysis proved more sensitive than the PSTHs based on timing alone (Fig. 1D): 61 out of 84 (73%) recorded Purkinje cells showed some degree of coherence (>0.5) between complex spike firing and licking (Fig. 4A). The complex spike activity of each individual Purkinje cell had a preferential phase within the licking cycle, but the preferred phases of the different cells were heterogeneously and relatively randomly distributed, so that together they covered the entire cycle, although a slight preference for the onset of protrusion and the maximal protrusion was observed (Fig. 4A). We next plotted the coherence levels for the complex spike modulations of each Purkinje cell based on the insertion points of the electrodes at the surface of the cerebellar hemispheres and vermis. This structure-function analysis did not reveal specific anatomical hotspots (Fig. 4B).

**Fig. 4.**
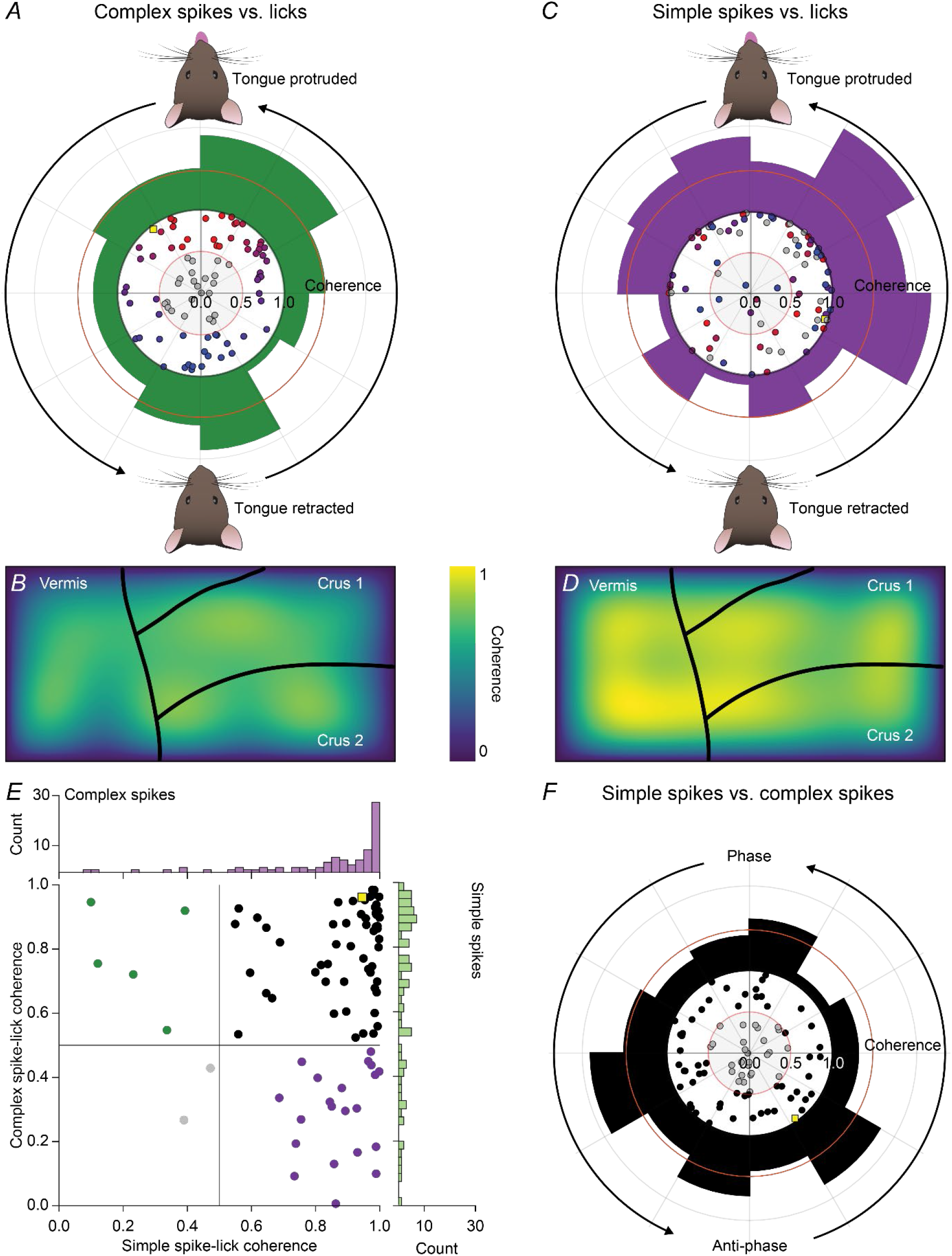
Coherent activity between complex spikes, simple spikes, and licking. **A** - Complex spike distributions around licks show high coherence values (>0.5) in 27/50 cells. The preferential phase in which complex spikes occur in those cells varies along the cycle. Each dot represents a cell and is colour-coded by the distance of its preferential phase during rhythmic tongue movements. The yellow square represents the same example cell shown in Fig. 1 (notice that high coherence value and location in the third quadrant reflect the oscillating behaviour and a main peak that follows the tongue detection respectively, as observed for the complex spike PSTH from Fig. 1D). Outside: histogram showing the distribution along the lick cycle of cells with lick-complex spike coherence >0.5. **B** - Heatmap showing the average coherence level for complex spikes in the recording area based on the point of insertion of the recording electrodes on the surface of the cerebellum. **C** - Same plot as in A, but for simple spikes. **D** - Same plot as in B, but for simple spikes. **E** - Scatter plot showing the lack of a correlation between complex spike-lick coherence and simple spike-lick coherence, independent from the preferred phase. Some cells show only complex spike-lick coherence (green), others only simple spike-lick coherence (purple). **F** - When relating complex spike and simple spike modulation to licking, in 23/50 cells only the coherence values are above 0.5 and in general are lower than the ones observed between simple spikes and licking. A certain degree of antiphase becomes visible.

The coherence analysis showed that the phase relation between simple spike firing and licking was stronger than that for the complex spikes (Fig. 4C). In total 77 out of 84 (92%) Purkinje cells showed >0.5 coherence with licking. Moreover, when we spatially plotted the coherence levels for the simple spike modulations, specific anatomical hotspots emerged, with the strongest levels of coherences localized at the border regions between hemispheres and vermis (Fig. 4D).

This analysis revealed that most Purkinje cells (56 out of 84; 67%) showed both coherence between complex spikes and licking, as well as between simple spikes and licking (Fig. 4E). Instead, just 7% (6 out of 84) of the Purkinje cells displayed coherence with the licking for their complex spikes only, and 24% (20 out of 84) for simple spikes only (leaving only 2 cells with no clear coherence). The phase relations between complex spikes and simple spikes varied strongly between Purkinje cells. In general, though, the majority of Purkinje cells had a tendency to fire complex spikes and simple spikes in antiphase (Fig. 4F).

*Are Purkinje cells also rhythmically active at rest?* The presence of coherence between licking and spiking substantiates the presence of rhythmic firing during licking bouts. Given that Purkinje cells are often involved in different types of behaviour (Cao *et al*., 2012; Romano *et al*., 2020), and that orofacial behaviours typically show similar dynamics (Moore *et al*., 2013; Romano *et al*., 2020), we wondered whether rhythmic Purkinje cell activity could also be found during periods without licking activity. Hence, we constructed auto-correlograms of simple spike activity during and in between licking bouts. This analysis confirmed rhythmic simple spike activity during licking, but little to no rhythmic activity in the absence of licking (power of simple spike autocorrelograms at licking frequency: during lick bouts: 2.7 (1.7) vs. in between lick bouts: 1.4 (0.7), p < 0.001, paired t test, r = 0.68, p < 0.001, Spearman rank correlation, Fig. 5). Thus, we conclude that the occurrence of rhythmic simple spike firing was largely confined to periods with licking.

**Fig. 5.**
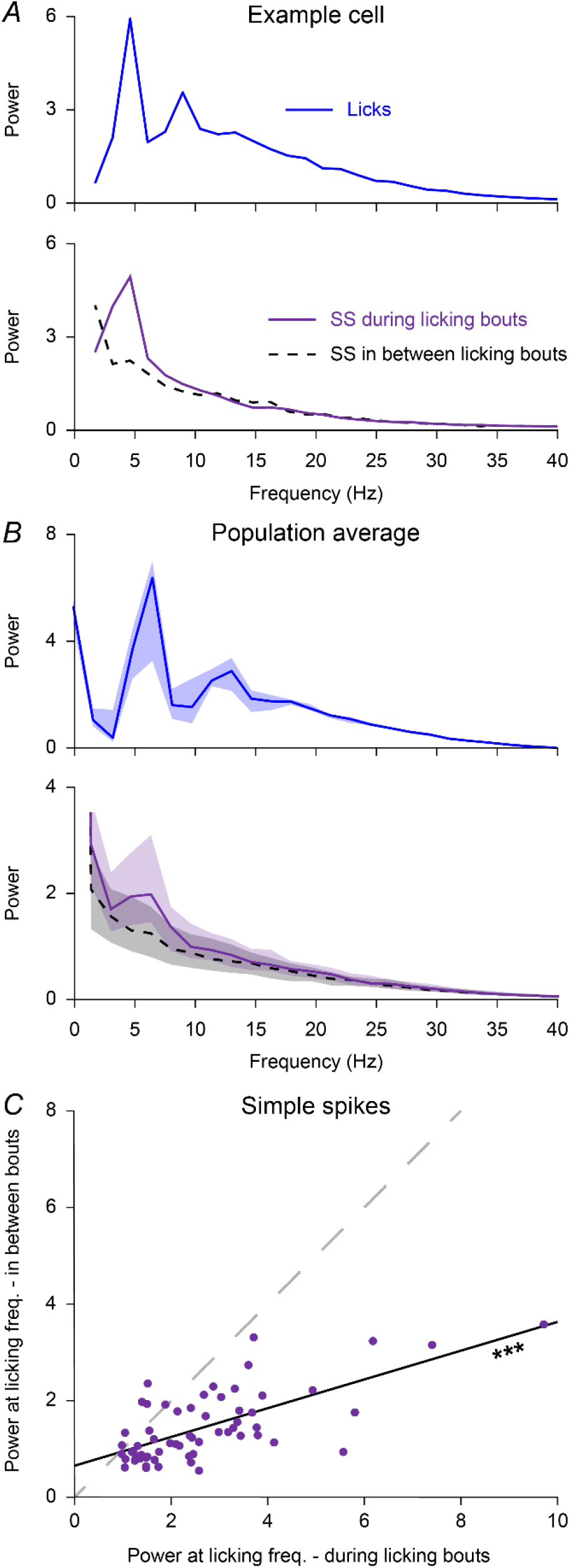
Oscillations in phase with licking are absent outside of licking bouts. **A** - Top: power spectrum calculated from the autocorrelogram of licks during a randomly selected session of spontaneous licking. During this session, licking was rhythmic at a frequency of approximately 5 Hz. Bottom: rhythmic firing at the same frequency was also evident in the autocorrelogram of simple spikes (SS) fired during, but not in between, licking bouts. **B** - Power spectra of the autocorrelograms of licking (top) and of simple spike firing (bottom) for the 61 out of 84 recorded Purkinje cells that showed an evident peak around the licking frequency. Medians with shades representing inter-quartile range (IQR). **C** - For every of the 61 Purkinje cell included in the analysis of panel B, the power at the preferred licking frequency during that recording during vs. in between licking bouts. The power was typically stronger during licking bouts (p < 0.001, Wilcoxon matched-pairs test), yet correlated per Purkinje cell (r = 0.606, p < 0.001, Spearman rank correlation).

### Purkinje cell activity correlates with the trajectory of the tongue

After establishing that Purkinje cell simple spike firing can be strongly linked to rhythmic licking, we wondered to what extent simple spike firing affects tongue movements. To this end, we analysed video recordings of spontaneous licking to monitor the variation in tongue movements between trials. For each mouse, we plotted the locations of maximal protrusion of each lick, and divided the distribution of end-points in tertiles, both on the rostro-caudal and on the left-right axis (Fig. 6A). For each one of the nine resulting areas, we looked at the average normalized simple spike firing frequency preceding the end point of each lick at two time intervals, the first one 150 to 75 ms before and the second one 75 to 0 ms before the moment of maximal tongue protrusion (Fig. 6B).

**Fig. 6.**
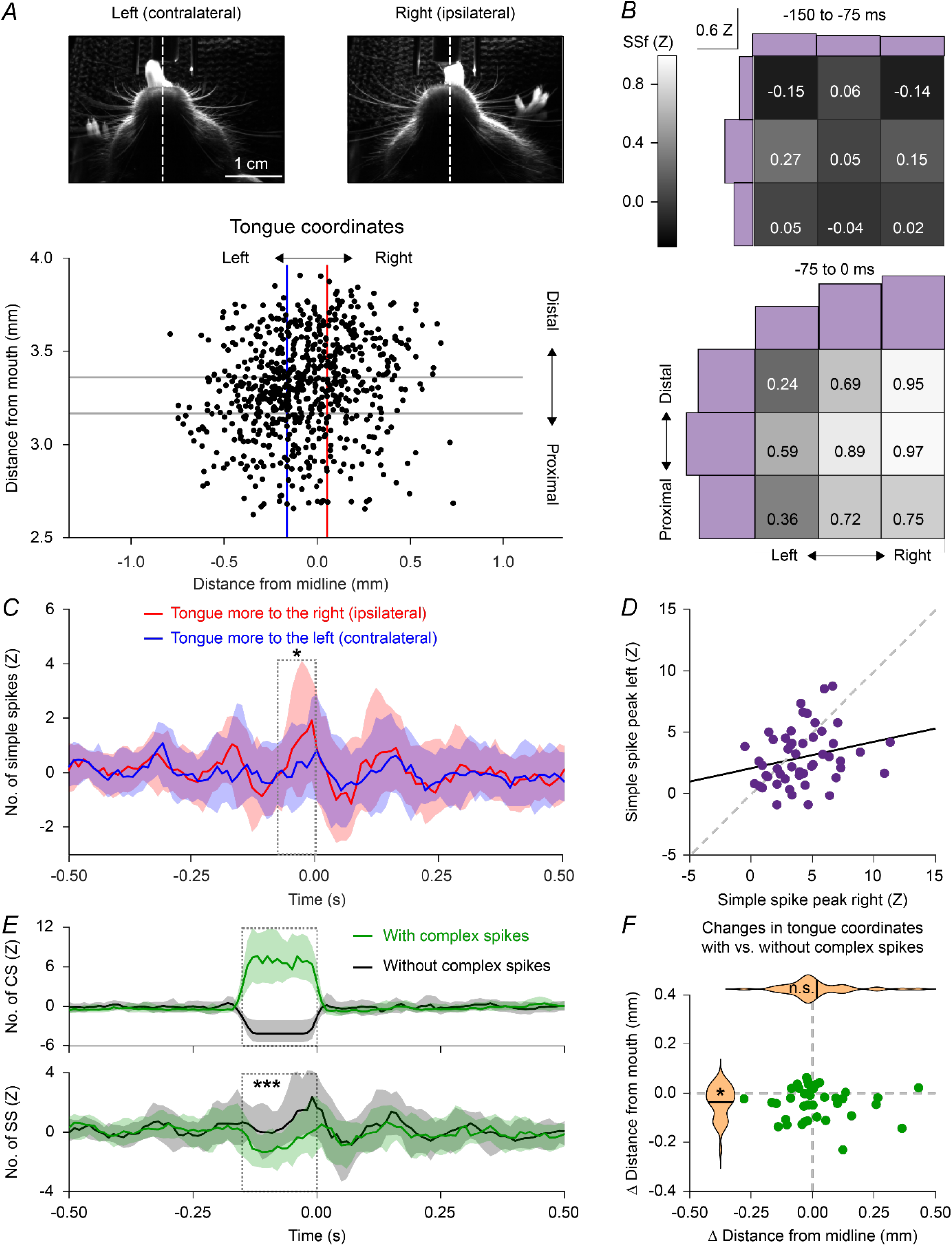
Purkinje cells encode the coordinates of tongue protrusion. A - Using video analysis, licks were classified based on the coordinates of the tongue at the moment of maximal protrusion. Two example frames for licks to the left and to the right are shown. For the whole session, the rosette resulting from the tongue coordinates is divided in tertiles on the distal-proximal and left-right axes. **B** - Heatmaps showing the average simple spike frequency (in Z scores) preceding licks in two consecutive time intervals relative to the moment of maximal protrusion (top and bottom) for the nine spots obtained after the subdivision of each rosette. Histograms of the spike frequencies along the two axes are shown in purple. **C -** Median simple spike PSTHs showing increased simple spike firing during tongue protraction (between 75 and 0 ms prior to reaching the maximal protraction) of licks bended ipsilaterally vs. contralaterally: p = 0.021, n = 84 Purkinje cells, paired t test. Medians with shades representing IQR. **D -** Scatter plot of the peaks in simple spike firing during the interval highlighted in C. r = 0.160, p = 0.262, n = 84 Purkinje cells, Spearman rank correlation. **E** - PSTHs showing the distribution of complex spikes (top) and simple spikes (bottom) for licks where complex spikes occurred or not in a time window of 75 ms preceding each lick. *** p = 0.001, n = 84 Purkinje cells, paired t test. Medians with shades representing IQR. **F** - Tongue protrusion of licks preceded by complex spikes was shorter, but did not have a lateral bias (violin plots: black lines indicate mean). Tongue extension (y axis): p = 0.001; bending (x axis) p = 0.431, n = 84 Purkinje cells, paired t tests).

Next, we quantified the asymmetry in the simple spike firing rate during tongue protrusion (75 to 0 ms before reaching the maximal protrusion) of licks inclined to the right or the left side. More simple spikes were fired on average in this 75 ms interval during right-bended (ipsilateral) licks (mean simple spike frequency (Z): ipsilateral 1.24 (2.54) vs. contralateral 0.34 (2.22), p = 0.021, paired t test, Fig. 6C). Analogously, the peak values of the two distributions also differed (peak simple spike frequency (Z): ipsilateral 4.01 (2.52) vs. contralateral 2.94 (2.21), p = 0.013, paired t-test). The contrast between ipsi- and contralateral licks was emphasized by the absence of a significant correlation in maximal simple spike firing during both conditions (r = 0.160, p = 0.262, Spearman rank correlation, Fig. 6D). This suggests that simple spikes may enhance tongue movements towards the ipsilateral side.

We next investigated whether complex spikes could also play a role in the laterality of tongue movements, as complex spikes are followed by a pause in simple spike firing. Trials with a complex spike display less simple spikes (mean Z-score of the 150 ms interval before lick detection: without CS: 0.92 (1.40); with CS -0.69 (1.37), p < 0.001, paired t-test. Fig. 6E). This did not result in a lateral bias of tongue movements (- 0.01 (0.27) vs. 0.01 (0.29) mm; p = 0.431, paired t test), but rather in a reduction in the extension of tongue protrusion (4.18 (0.88) vs. 4.15 (0.89) mm; p = 0.001, paired t test, Fig. 6F).

### Adaptation of Purkinje cell activity during targeted tongue movements

Next, we tested the ability of mice to adapt their tongue movements to a displaced target (Fig. 7A). To this end, we moved the lick-port 3 mm to the right during the retraction phase of randomly selected licks. The lick-port stayed at that position during 750 ms and then returned to the central position. While the first lick after lick-port displacement did not yet show statistically significant bending, the second and third licks were adapted to the new target position (p < 0.001, Friedman’s ANOVA with post-hoc tests (based on tongue position in mm) vs. lick 0: lick 1: p = 0.568, lick 2: p = 0.002, lick 3: p < 0.001, n = 13 mice, Fig. 7B, C). The movement back to the centre had very similar impact (p < 0.001, Friedman’s ANOVA with post-hoc tests vs. lick 0: lick 1: p = 0.772, lick 2: p < 0.001, lick 3: p < 0.001, n = 13 mice, Fig. 7C).

**Fig. 7.**
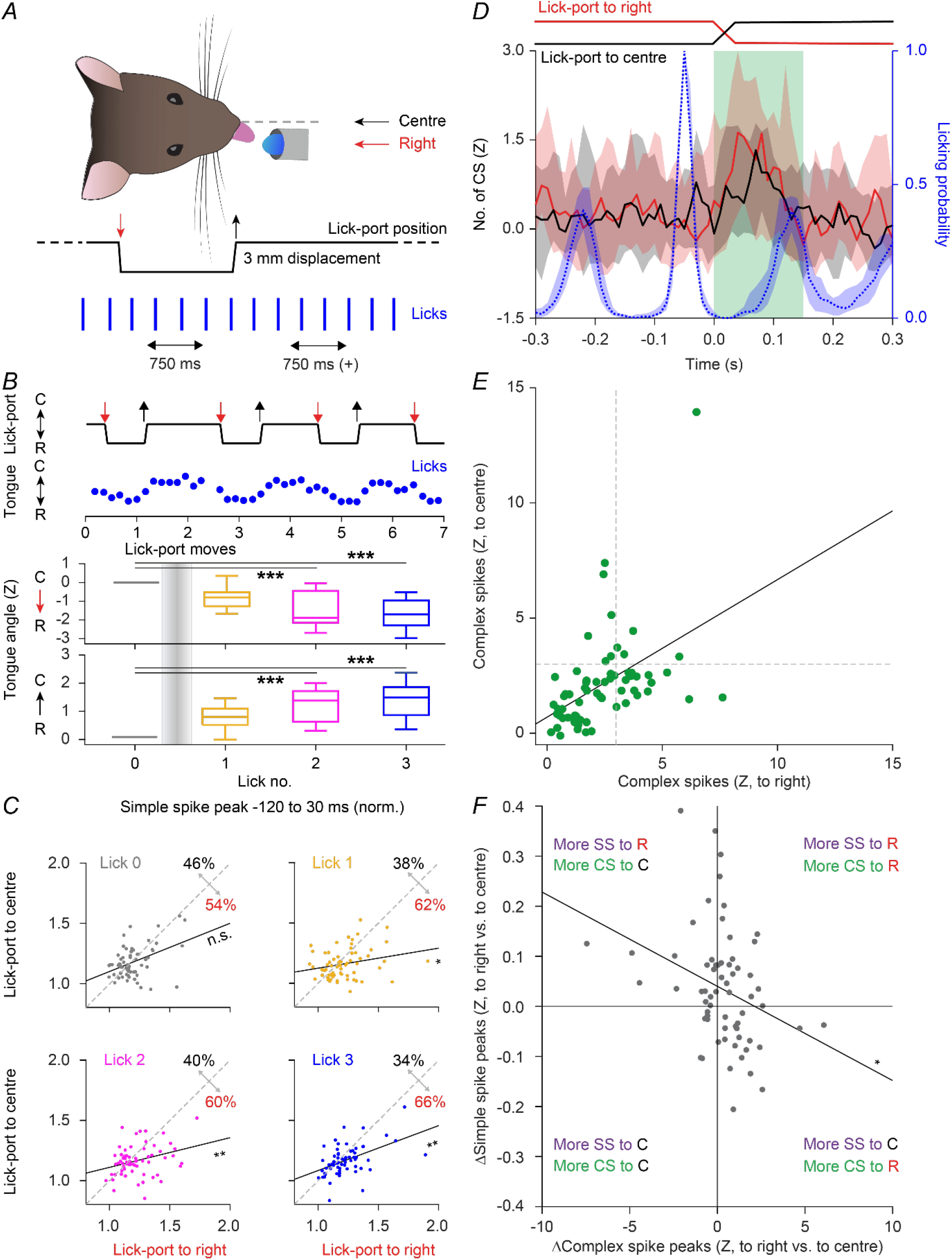
Purkinje cell firing reflects motor adaptation. **A** - Experimental scheme: 3 mm rightward movements of the lick-port occurring during the retraction phase of the tongue were triggered on randomly selected licks. The lick-port was moved back to the original position after 750 ms and staid in the central position for a minimum of 750 ms, so that the direction of the 4 to 5 licks following each movement needed to be adjusted for a more efficient consumption of the water. Outside of the 1.5 s trial time, some licks were isolated from any lick-port movement for a minimum of 300 ms and used as baseline. **B** - A randomly selected epoch showing that left-right changes in position of the lick- port lead to left-right changes in the endpoints of tongue protrusions. Mice (n = 13) adapted the direction of the tongue protrusions partially for the first lick following the trials, fully for the second and third licks. p < 0.001, Friedman’s ANOVA, with post- hoc tests comparing the most extended position (in mm) of the tongue between licks 1-3 vs. lick 0 (rightward movements: lick 1: p = 0.568, lick 2: p < 0.001, lick 3: p < 0.001; centreward movements: lick 1: p = 0.772, lick 2: p < 0.001, lick 3: p < 0.001). **C** - Scatterplots of the peak values of the simple spike distributions around licks of interest for lick-port movements to the right and back (rightward vs. centreward movements, lick 0: p = 0.235, lick 1: p = 0.018, lick 2: p = 0.003, lick 3: p = 0.002, paired t- test). **D** - PSTHs showing complex spike responses elicited after right-ward (red) or centre-ward (black) movements of the lick-port (27 / 65 cells with a peak response larger than 3 SD). Complex spike responses typically preceded the moment of maximum protrusion of the tongue of the subsequent lick (see lick distribution in blue). Medians with shades representing IQR. **E** - Correlation between the peak complex spike responses elicited by lick-port movements (r = 0.46, p < 0.001, Spearman correlation). **F** - Correlation between the difference in selectivity (right - centre) for complex spike peaks and the average simple spike peaks around licks 2 and 3 (r = -0.32, p = 0.010, Spearman correlation).

Lick-port movement triggered increased complex firing in 27 (42%) out of 65 Purkinje cells (Fig. 7D). There appeared to be little direction-selectivity in this complex spike response, as the peaks of complex spike modulation to the lick-port moving rightward or back to centre were positively correlated (r = 0.46, p < 0.001, Spearman correlation, Fig. 7E).

In line with the finding that simple spike firing encodes spontaneous fluctuations in the laterality of tongue movements (Fig. 6B-D), we observed changes in simple spike firing prior to consecutive licks following the movements of the target that reflected the changes in the direction of tongue protrusions (SS peak normalized to lick 0: lick 1, to right 1.21 (0.18) vs. to centre 1.15 (0.12), p = 0.018, lick 2, to right 1.23 (0.16) vs. to centre 1.16 (0.13), p = 0.003, lick 3, to right 1.22 (0.16) vs. to centre 1.16 (0.12), p = 0.002, paired t-test, Fig. 7F). Overall, the complex spike responses were anti-correlated with the changes in simple spikes (r = -0.32, p = 0.010, Spearman correlation. Fig. 7G). Thus, complex spikes signalled the sensory event of the moving lick-port, while the simple spikes correlated with the trajectory of the altered tongue movements.

### Purkinje cell stimulation results in bending of the tongue

Increased simple spike firing correlated with ipsilateral bending of the tongue during protrusion. To find out if this may be a causal relation, we used optogenetic stimulation of Purkinje cells to evoke increased simple spike firing (Witter *et al*., 2013; Lindeman *et al*., 2021). We used randomly selected licks to trigger optogenetic stimulation in phase with licking. An optic fibre of 400 μm diameter was placed on the surface of the cerebellum at the border between crus 1 and crus 2, just right of the vermis (Fig. 8A), as this is the area with the strongest coherence between simple spike activity and licking (Fig. 4D). This stimulation resulted in a reduced lick probability of the consecutive lick when applied during the retraction phase of the previous lick in an example mouse (Fig. 8B).

**Fig. 8.**
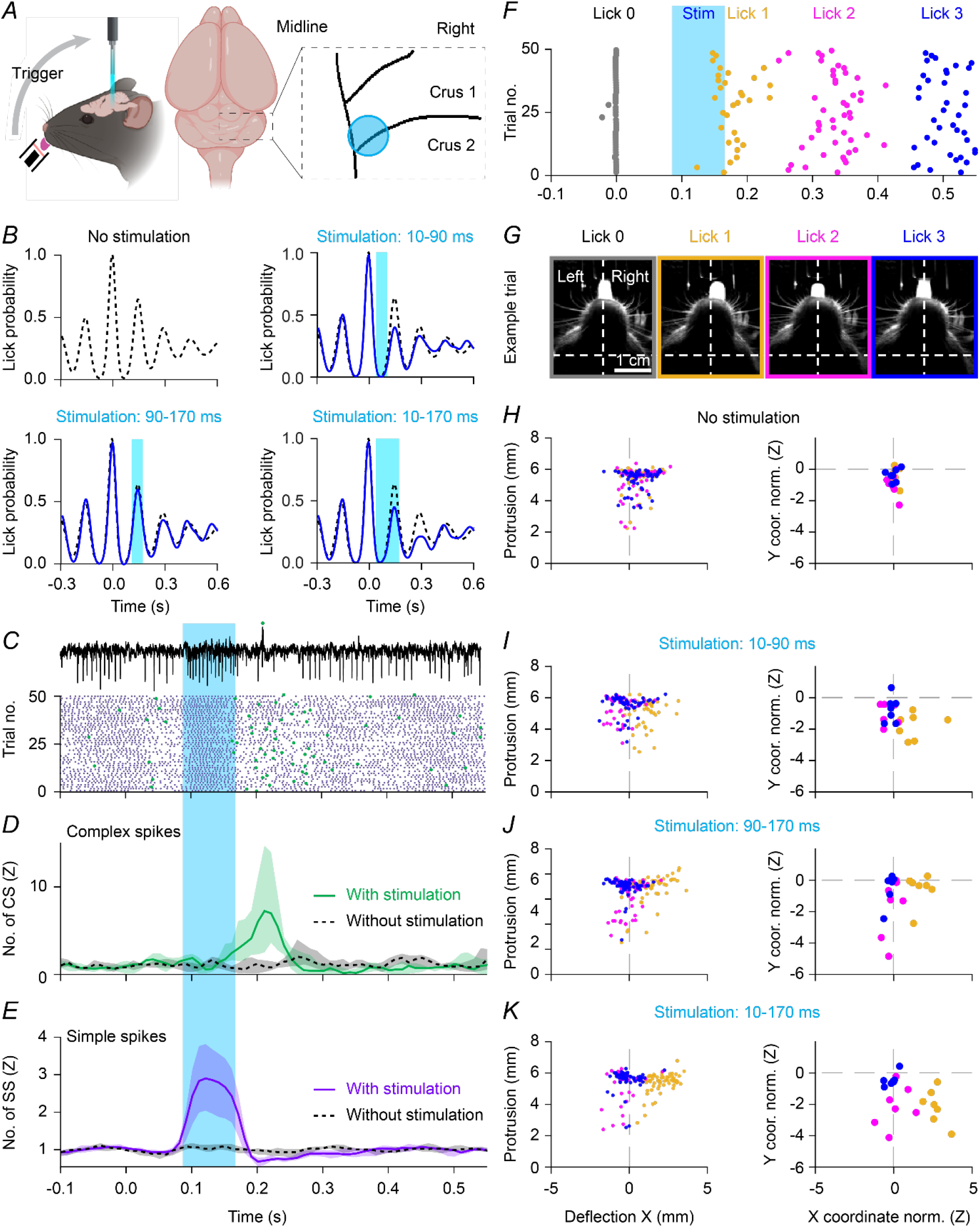
Optogenetic stimulation affects the execution of licking. **A** - Optogenetic stimulation was performed by shining blue light (470 nm) on Purkinje cells expressing ChR2 under control of the Pcp2 promotor. We used an optical fibre placed at the border of crus 1 and crus 2 close to the vermis on the right-hand side of the brain. Optogenetic stimuli were triggered by random licks, and we used three different light pulses either starting at 10 or 90 ms after lick detection and lasting 80 ms, or a longer pulse starting 10 ms after lick and lasting 160 ms. **B** - For an example mouse and for the three different optogenetic stimulation conditions, the distribution of licks that efficiently crossed the lick-port sensor (blue) vs. those in the absence of optogenetic stimulation (black). **C** - Example of an extracellular recording of a Purkinje cell during optogenetic simulation and a raster plot of complex spikes (green) and simple spikes (purple) distribution around light stimulations. Median with shade representing IQR. **D** - Complex spikes PSTHs for trials with and without optogenetic stimulation (n = 10 Purkinje cells). Median with shade representing IQR. **E** - As D, but for the simple spikes. **F** - Timing of occurrence of maximal tongue protrusions around optogenetic stimulations for an example mouse obtained after video tracking, for stimulation 90 to 170 ms after randomly selected licks (lick 0, grey). **G** - Example frames belonging to four consecutive licks of a trial during optogenetic stimulation of the right hemisphere. Notice that the first lick after stimulation is bended to the right, the second lick is shorted, the third lick is comparable to the one before stimulation. **H** - Left: for an example mouse, coordinates of licks 1, 2 and 3 (okra, magenta and blue) following randomly selected control licks not followed by optogenetic stimulation. Right: each datapoint is now representing the average value of individual mice (n = 7 mice). **I** - Same as in H, but for optogenetic stimulations 10 to 90 ms after detection of lick zero (right, x-coordinate lick 1 vs lick 0: p = 0.016, paired t tests with Bonferroni correction; n = 7 mice). **J** - Same as in H, but for optogenetic stimulations 90 to 170 ms after detection of lick zero (right, x-coordinate lick 1 vs lick 0: p < 0.001). **K** - Same as in H, but for optogenetic stimulations 10 to 170 ms after detection of lick zero (right, x-coordinate lick 1 vs lick 0: p < 0.001).

Further analysis revealed that optogenetic stimulation triggered a robust increase in simple spikes, which was followed by complex spike firing approximately 70 ms after the end of stimulation (Fig. 8C-E), as observed previously (Witter *et al*., 2013). We tested the impact of stimulation either during tongue retraction of the detected lick, tongue protrusion of the next lick, or using a twice as long interval (160 instead of 80 ms) to cover both. Optogenetic stimulation resulted in altered tongue movements: the first lick occurring after stimulation onset was bended ipsilaterally (x-coordinate lick 1 vs lick 0, 10-90 ms: p = 0.016; 90-170 ms: p < 0.001; 10-170 ms: p < 0.001, paired t tests with Bonferroni correction, n = 7 mice, Fig. 8F-K, okra). Thus, optogenetic stimulation of Purkinje cells caused ipsilateral bending of the tongue, suggesting that the simple spike rate indeed affects the tongue trajectory.

In another batch of mice, we introduced a second optic fibre at the opposite hemisphere (Fig. 9A), and stimulated in random fashion either one side at a time or both simultaneously, using the longer stimulus interval of the previous experiment (10-170 ms after lick detection). Stimulating on the right side induced, as before, a right-ward bending of the next tongue protrusion (lick 1 right stim. vs lick 1 no stim, p = 0.029, Wilcoxon rank-sum test, n = 4 mice), while stimulating on the left caused a left-ward bending (lick 1 left stim. vs lick 1 no stim, p = 0.029, Wilcoxon rank-sum test. Fig. 9C-D). Bilateral stimulation resulted in reduced extension of the tongue with no lateral bias (lick 1 with bilateral stim. vs lick 1 without stim, x-coordinate, p = 0.114, Wilcoxon rank-sum test), indicating that enhancing simple spike firing, thus inhibiting the cerebellar nuclei, could suppress the contraction of ipsilateral muscles. With a last group of mice, we investigated the impact of more lateral stimulation and found that they were significantly less effective in generating ipsilateral bias in tongue movements (lick 1 medial stim. vs lick 1 no stim, p = 0.048, lick 1 lateral stim. vs lick 1 no stim, p = 0.400, Wilcoxon rank-sum tests with Bonferroni correction, n = 4 mice. Fig. 9E). Thus, artificially increasing simple spike firing in phase with licking could affect the execution of individual cycles of tongue movements during licking, proving the causal relation between Purkinje cells output and directionality of tongue movements.

**Fig. 9.**
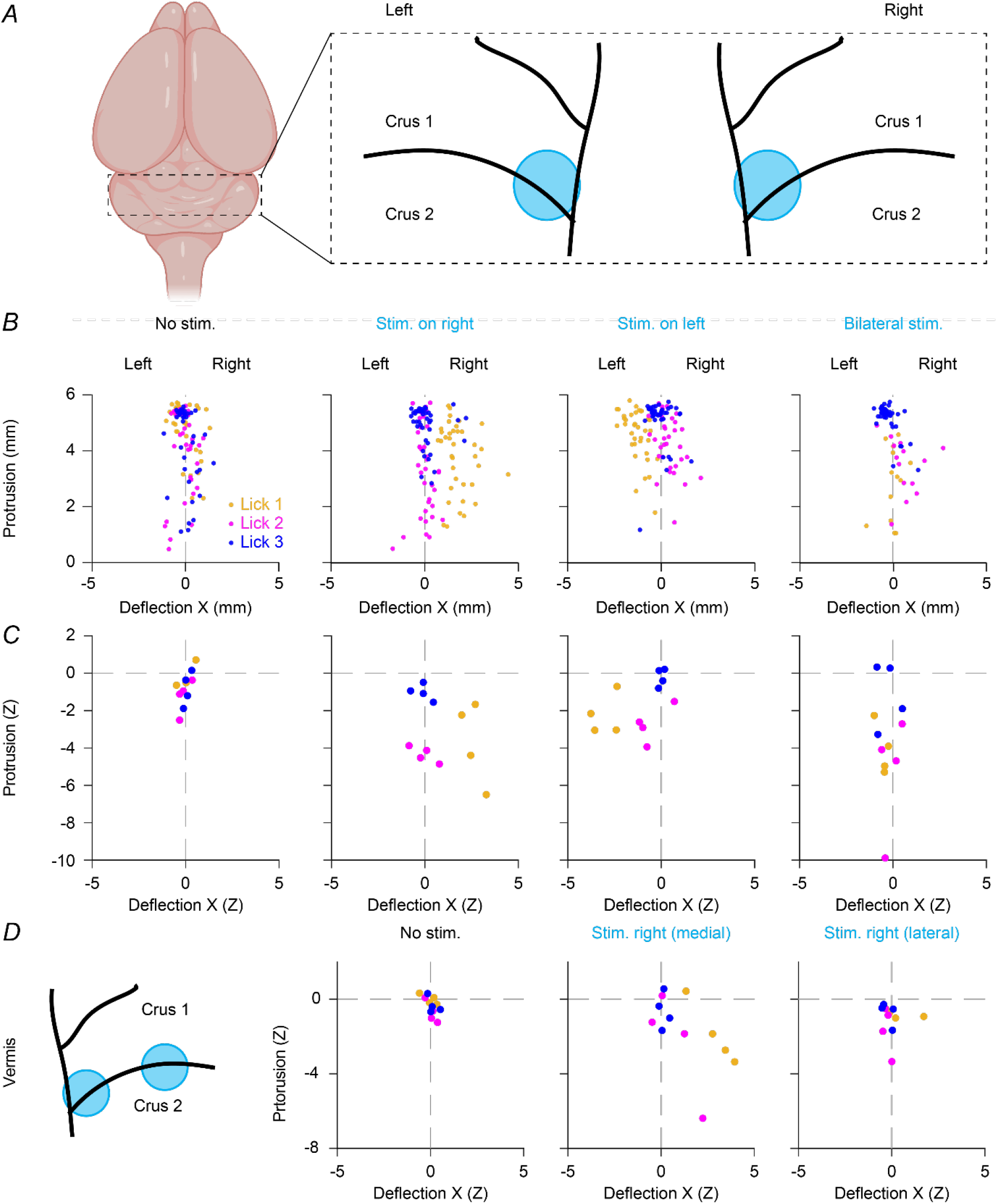
Stimulation of the medial hemispheres inhibit the ipsilateral extension of tongue muscles. **A** - Two optic fibres were placed on the right and on the left side of vermis, between lobules crus 1 and crus 2, and were activated either alone or simultaneously during licking. **B** - For an example mouse, coordinates of licks 1, 2 and 3 (okra, magenta and blue) in trials without optogenetic stimulation, with stimulations of the right hemisphere, with stimulations of the left one, and with simultaneous stimulations of both sides. **C** - Same as in B, but each datapoint is now representing the average per mouse (x-coordinate, lick 1 right stim. vs lick 1 no stim, p = 0.029; lick 1 left stim. vs lick 1 no stim, p = 0.029; lick 1 with bilateral stim. vs lick 1 no stim, p = 0.114, Wilcoxon rank-sum test; n = 4 mice). **D -** For a different set of four mice, one fibre was placed on the right side as for previous experiments, and a second fibre was placed more laterally (left). Lateral stimulations were less effective (lick 1 medial stim. vs lick 1 no stim, p = 0.048, lick 1 lateral stim. vs lick 1 no stim, p = 0.400, Wilcoxon rank-sum tests with Bonferroni correction.

## Discussion

Mice, like most mammals, make fast rhythmic tongue movements when drinking. It is generally assumed that this rhythm is set by a central pattern generator in the brainstem (Brozek *et al*., 1996; Travers *et al*., 1997; Dempsey *et al*., 2021; Kleinfeld *et al*., 2023). Notably, also groups of cerebellar Purkinje cells turn out to modulate their firing pattern in such a way that they encode specific aspects of tongue movement. Complex spikes can signal unexpected changes in the target position of the tongue and simple spikes are associated with the (a)symmetry of tongue movements. By challenging mice to adjust their tongue movements towards a quickly moving target for licking, we could demonstrate that alterations in simple spike firing are indeed consistent with altered movements. Finally, optogenetic stimulation of Purkinje cell activity results in changes in the tongue trajectory. Thus, the activity of Purkinje cells does not only reflect the way goal-directed tongue movements are executed, but it can actually also alter these.

### Tongue movements

Mammalian tongues thank their motility to a combination of four extrinsic and four intrinsic muscles, and most of these muscles are paired. Extrinsic muscles connect the tongue with surrounding bones, and intrinsic muscles are located entirely within the tongue. One could consider the genioglossus muscle as the driving force of tongue protrusion, and the hyoglossus muscle the main actor of tongue retraction, but tongue movements are typically the result of simultaneous activity of multiple tongue muscles (Bennett, 1937). The initial conclusion that left-or rightward bending of the tongue results solely from unilateral activation of the genioglossus muscle (Braus, 1924), is therefore likely an oversimplification. In reality, deflections of the tongue are caused by the asymmetric activation of several tongue muscles (Bennett, 1937; Abd-El-Malek, 1938; Lowe, 1980).

Seven out of eight tongue muscles are controlled by motor neurons in the hypoglossal nucleus. Only the (extrinsic) palatoglossus muscle is innervated from the vagus nerve. Accordingly, the hypoglossal motor neurons produce rhythmic activity in tune with tongue movements (Wiesenfeld *et al*., 1977). A plausible origin of this rhythmic activity is a group of *Phox2b*-positive pre-motor neurons in the intermediate reticular formation (Dempsey *et al*., 2021), which is the brain area supplying most of the afferent fibres to the hypoglossal nucleus (Guo *et al*., 2020). In itself, the brainstem circuit is likely to be sufficient for the generation of reflexive rhythmic tongue movements, but it does require input from the cerebral cortex for volitional tongue movements (Bignall & Schramm, 1974; Grill & Norgren, 1978; Bollu *et al*., 2021; Kleinfeld *et al*., 2023).

### Cerebellar activity during licking

In our study, complex spikes were found to encode the phase of rhythmic tongue movements, in line with previous work in rats (Welsh *et al*., 1995). We observed such complex spike modulation in markedly fewer Purkinje cells than simple spike modulation, and this may explain why two previous studies failed to measure complex spike modulation during rhythmic licking in mice (Bryant *et al*., 2010; Cao *et al*., 2012). In contrast to the relatively weak encoding of individual licks by complex spikes, we observed a much larger relation between behavioural state changes, thus the start or end of licking bouts, and complex spike firing.

Complex spike firing just prior to tongue protrusion generally results in slightly shorter tongue extensions, pointing towards a role of complex spike firing in motor control. Complex spikes just prior to movement have previously been correlated with alteration in movement kinetics of limbs and whiskers (Kitazawa *et al*., 1998; De Gruijl *et al*., 2014; Romano *et al*., 2018), and we show here that this may apply to tongue movements during state changes as well.

Although we observed a potential impact of complex spike firing on tongue movements during state changes, it is assumed that it are mainly the simple spikes that affect motor execution in general (Chen *et al*., 2016; Bina *et al*., 2021). According to prevailing theories of cerebellar function, complex spikes can serve to encode sensory prediction errors. They translate these to cerebellar learning by regulating the synaptic plasticity of parallel fibre inputs and thereby simple spike firing patterns (Marr, 1969; Albus, 1971; Ito, 2000; Yang & Lisberger, 2014; Romano *et al*., 2018). Here, we induced sensory prediction errors by suddenly moving the target for licking. This unexpected event triggered robust complex spike firing, followed by changes in simple spike firing and alterations in the trajectories of the tongue.

During licking, rhythmic simple spike firing was observed abundantly in the lateral cerebellum, in line with previous results (Bryant *et al*., 2010; Cao *et al*., 2012; Gaffield *et al*., 2022). We also found that the majority of cells that modulate rhythmically are located more medially within the hemispheres. Each Purkinje cell has a preferred phase lag compared to tongue movement, so that together Purkinje cells cover the whole lick cycle, similar to other rhythmic behaviours like walking (Sauerbrei *et al*., 2015) and respiration (Romano *et al*., 2020). As during respiration (Romano *et al*., 2020), there is an uneven distribution of preferred phases of individual Purkinje cells over the licking cycle. Relatively many Purkinje cells are preferably active during the later phases of tongue protrusion. Simple spike firing during this period correlates with an ipsilateral bending of the tongue. The Purkinje cell output controls the activity in the cerebellar nuclei, which in turn modify activity of the neurons downstream in the reticular formation that are engaged in licking (Lu *et al*., 2013).

*The cerebellum and the symmetry of movements* Healthy humans walking on a split-belt treadmill can easily adapt their stepping pattern to considerable speed differences between their right and left side. Patients with cerebellar damage, however, have trouble to generate predictive compensation mechanisms (Morton & Bastian, 2006; Hoogkamer *et al*., 2015). A similar inability to create asymmetric gaiting patterns was observed in cerebellar mutant mice (Darmohray *et al*., 2019). Also for whisker movements, which can show asymmetry to compensate for head movements (Towal & Hartmann, 2006), cerebellar activity is required to generate asymmetry. In particular, complex spikes can alter the degree of symmetry between whisker movements on either side of the head (Romano *et al*., 2022). Given that changes in complex spike firing can be correlated to the (a)symmetry of tongue movements due to their suppressing effect on Purkinje cell simple spike output, we propose that the cerebellum is crucial for creating various types of asymmetric movements and that the cerebellar mechanisms engaged to create the asymmetry depend on the specific structure of the motor plant involved as well as the synaptic polarity of the intermediate neuronal hubs (De Zeeuw, 2021). For tongue movements, we found a strong impact of increases in simple spike firing on determining the angle of individual licks within a licking bout.

### Anatomical projections from the cerebellum to the hypoglossal nucleus

There are multiple pathways along which the cerebellum can affect the execution of tongue movements. First of all, there is a direct but sparse projection from the cerebellar nuclei to the hypoglossal nucleus (Guo *et al*., 2020; Judd *et al*., 2021; Novello *et al*., 2024). Indirect projections between the cerebellum and the hypoglossal nucleus may be numerically and thereby functionally more relevant, as in other motor systems (Novello *et al*., 2024). Importantly, the cerebellum provides many of the non-brainstem inputs to the *Phox2b*-positive pre-motor neurons in the intermediate reticular formation (Dempsey *et al*., 2021), making this cluster of neurons that is tentatively considered a central pattern generator for tongue movements, a likely intermediate for cerebellar output. Given that the cerebellum broadly projects to brainstem regions (Teune *et al*., 2000; Novello *et al*., 2024), and that many, if not most, of these project to the hypoglossal nucleus (Guo *et al*., 2020), also other brainstem-mediated indirect pathways may serve to convey cerebellar output to the hypoglossal nucleus. Finally, the cerebellum could also modulate tongue movements indirectly via the thalamus and tongue motor cortex (Aoki *et al*., 2019).

### Clinical relevance

The inability to control the tongue properly can lead to dysarthria, dysphagia and respiratory problems, which, in turn, can have serious impact on the quality of life, and, via aspiration pneumonia, even be lethal (Daniels *et al*., 1999; Takizawa *et al*., 2016; Duan *et al*., 2020; Krohn *et al*., 2023). There are multiple possible causes for oral movement disorders, of which impairment of the basal ganglia, as in Parkinson’s and Huntington’s disease, is the best characterized (Leopold & Kagel, 1996; Reilmann *et al*., 2010; Minagi *et al*., 2018). Neurodegenerative diseases and stroke are among the diverse neurological conditions that can impair tongue function, and thereby cause problems with speech and ingestion (Tao *et al*., 2012; Takizawa *et al*., 2016). Notably, both anterior and posterior circulation infarcts can affect tongue motility (Tao *et al*., 2012). The impact of anterior circulation infarcts can probably largely be explained by the impact of cerebral motor areas, especially the facial area of the primary motor cortex, which projects directly to the hypoglossal nucleus in humans and other primates (Kuypers, 1958b, a; Jürgens & Alipour, 2002; Morecraft *et al*., 2014). In addition, there are also indirect projections from the motor cortices to the hypoglossal nucleus, possibly via the *Phox2b*-positive neurons of the intermediate reticular formation (Jürgens & Alipour, 2002; Dempsey *et al*., 2021). While the direct cortico-hypoglossal projections seem to be limited to primates, the indirect projections are also abundant in other mammals (Jürgens & Alipour, 2002; Dempsey *et al*., 2021).

Damage to the cerebellum could explain, at least in part, the impact of the posterior circulation infarcts on the ability to control the tongue. Accordingly, problems with speech and swallowing are prevalent in patients with diverse forms of cerebellar ataxia (Tan *et al*., 2000; Ikeda *et al*., 2012; Ushe & Perlmutter, 2012; Markovic *et al*., 2016; Keage *et al*., 2017; Rezende Filho *et al*., 2019; Woo *et al*., 2019; Giardina *et al*., 2020). Functional brain imaging data are in line with the activation of the cerebellum during various tongue-related behaviours (Corfield *et al*., 1999; Dimitrova *et al*., 2006; Grabski *et al*., 2012; Ogura *et al*., 2012; Boillat *et al*., 2020; Groenendijk *et al*., 2020; Sörös *et al*., 2020). Likewise, cerebellar dysfunction can also be associated with hyperkinetic tongue movements (Salari *et al*., 2023). Tongue fasciculations, tremor and dystonia may occur in different forms of cerebellar ataxia, but predominantly in individuals that have also extracerebellar brain damage (Izumi *et al*., 2013; Salari *et al*., 2023). Thus, clinical observations imply the cerebellum in the control of the tongue, but not always as the primary cause of deficits in tongue movements.

The latter is in line with our observations. In general, similar to other behaviours (De Zeeuw *et al*., 2011), we failed to produce evidence that the cerebellum can initiate tongue movements. We found, however, that the firing pattern of cerebellar Purkinje cells can reflect specific aspects of tongue movement, including the length and the angle of protrusion. Both our data on the timing of Purkinje cell activity and on the impact of optogenetic stimulation suggest that Purkinje cells can adapt tongue movements.

### Conclusions

Tongue movements can be complex, and precise control of the tongue is required for such important processes as ingestion, respiration and speech. Reflexive tongue movements, triggered by sensory stimulation, can be coordinated solely by the brainstem and rely heavily on a central pattern generator in the intermediate reticular formation. Volitional control of the tongue requires the primary motor cortex, probably in conjunction with accessory motor areas of the cerebral cortex. Instead, the cerebellum is essential for fine motor control and adaptation to the behavioural context. Disruption of any of these regions leads to impairments in tongue control that can eventually be lethal.

## Competing interests

The authors declare they have no competing interests.

## Author contributions

Conception or design of the work: L.B., C.C., S.Y., L.W.J.B. and C.I.D.Z. Acquisition and analysis of data: L.B., C.C., S.Y., X.W. and L.W.J.B. Interpretation of data: L.B., C.C., S.Y., L.W.J.B. and C.I.D.Z. Drafting and revising article: L.B., S.Y., X.W., L.W.J.B. and C.I.D.Z. All authors have approved the final version of the manuscript and agree to be accountable for all aspects of the work in ensuring that questions related to the accuracy or integrity of any part of the work are appropriately investigated and resolved. All persons designated as authors qualify for authorship, and all those who qualify for authorship are listed above.

## Funding

This study was supported by Health-Holland to promote public-private partnerships (TKI-LSH EMCLSH21017: L.W.J.B.). C.I.D.Z. received financial funding from the European Union’s Horizon 2020 research and innovation program under the Marie Skłodowska-Curie grant agreement (#722098), Medical NeuroDelta Programme, Topsector Life Sciences & Health (Innovative Neurotechnology for Society or INTENSE), Albinism Vriendenfonds Netherlands Institute for Neuroscience, and European Research Council – Advanced Grant (#294775), NWO-Gravitation Program (DBI2).

## Acknowledgements

The authors thank Stéphanie Dijkhuizen and Nathalie van Wingerden for excellent technical support.

## Notes

### Competing Interest Statement

The authors have declared no competing interest.

